# Mechanical cues from muscle contraction regulate TGFβ signaling and epitenon formation during embryonic tendon development

**DOI:** 10.64898/2026.05.14.725162

**Authors:** Emily R. King, Leonardo Campos, Joanna Smeeton, Nadeen O. Chahine, Alice H. Huang

## Abstract

Muscle loading is required for embryonic tendon growth; however, the underlying mechanisms that regulate tendon development downstream of mechanical cues remain unidentified. Although tendons in muscle paralysis models are structurally and functionally inferior, whether these differences arise from cell or matrix deficits remains unclear. Analysis of *muscular dysgenesis* embryos by atomic force microscopy showed that structural and functional deficits in paralyzed tendon arise in part from reduced proliferation and collagen fibril disorganization. Bulk and single cell transcriptional analyses reveal that both collagenous and non-collagenous extracellular matrix components, as well as cytoskeletal and actomyosin-associated proteins, are dysregulated in *mdg* tendons, whereas tendon markers remain unchanged. Surprisingly, we find that an arrest of TGFβ signaling occurs during normal embryonic tendon growth and that TGFβ signaling is abnormally prolonged in paralyzed embryos. We also show for the first time, that specification of the epitenon depends on muscle contraction. Together, these findings establish cell and molecular requirements for muscle contraction in embryonic tendon development.

**Teaser:** Muscle contraction is required for embryonic tendon development through regulation of TGFβ signaling, epitenon formation, and matrix organization.

## Introduction

Tendons are dense connective tissues that transmit the forces generated by muscle contraction to bones to enable movement. Tendon collagenous matrix is synthesized and remodeled throughout development by resident tendon cells (*1, 2*). The dense, highly aligned matrix is composed primarily of type I collagen and is essential for enabling tendons to function effectively as tissue integrators and transmitters of mechanical load within the musculoskeletal system (*3, 4*). Other collagen types (III, V, XII), as well as non-collagenous components such as elastin and proteoglycans, are also implicated in fibril organization and regulation (*1, 5*). Surrounding the main tendon body is the epitenon, a thin connective tissue layer that is distinct from the main tendon body. The epitenon is predominantly composed of collagen IV and laminin, which together form a dense fibrillar network of unaligned fibers around tendons (*6, 7*). The epitenon not only encapsulates the collagen fibrils within the tendon proper to support gliding functions, but also contains the vascular and neural elements responsible for supplying blood flow and innervation to the tendon (*8, 9*). Tendons serve as essential integrating components of the musculoskeletal system, and their hierarchical organization is fundamental to maintaining normal physiological function.

*Scleraxis (Scx)* is the earliest marker for tendon cells during embryonic development and continues to label tendon cells through postnatal stages (*10, 11*). During mouse limb development, *Scx+* tendon progenitor cells are first detected at E10.5, and by E12.5 are aligned loosely between the adjacent muscle and skeletal structures (*11*). At E13.5, these tendon progenitors condense to form distinct tendon structures. This condensation occurs concurrently with progenitor cell differentiation, during which tendons begin to express additional tendon markers, such as *Tnmd, Mkx*, and *Egr1* (*12, 13*). Most of tendon patterning concludes by E14.5, after which embryonic tendon development proceeds mainly through continued elongation and growth of the tendons. During this period, tendons undergo lateral and longitudinal growth, likely due in part to tendon cell proliferation or progenitor cell recruitment or a combination of these processes (*14–16*). Coincident with the increasing size of tendons is the development of their functional properties through deposition and cross-linking of the tendon matrix (*17, 18*). The epitenon marker, *Tppp3*, is first detected in E13.5 limbs adjacent to tendons in the metacarpals and surrounding most tendons in the wrist. By E15.5, nearly all limb tendons are enclosed by epitenon (*19*). While *Tppp3* remains the only embryonic marker identified for epitenon cells, it is also detected in the developing muscles and surrounding some joints. The timing of *Tppp3* expression around tendons suggests that the formation of the epitenon subsequently follows the events of tendon differentiation (E13.5) and continues throughout the growth period (E14.5+). To date, there is almost nothing known about how the epitenon is specified or the regulators of epitenon development.

In previous studies, we and others showed a requirement for muscle contraction during the stages of embryonic tendon growth, following initial progenitor differentiation (*14, 18, 21, 22*). In chick embryos, rigid and flaccid muscle paralysis gives rise to significantly weaker and more disorganized tendons (*18, 22, 23*). In mouse embryos, the *muscular dysgenesis* (*mdg*) mutant has been a useful tool to study muscle paralysis. *Mdg* embryos harbor a spontaneous mutation that disrupts muscle excitation-contraction coupling, resulting in muscle paralysis (*24*). We previously showed that *mdg* tendons are grossly normal at E14.5 but are significantly thinner with minor patterning defects by E16.5 (*14*). Others have shown that the maturation of matrix at the myotendinous junction (muscle-tendon attachment) is also disrupted in *mdg* mutants (*25*). In *mdg* knees, tendon and ligament structures are also smaller with cell and matrix abnormalities (*26*). While the *mdg* tendons are smaller, it is unclear whether this is due to deficits in cells or matrix, and the molecular basis of these changes is also unclear.

In the present study, we determine how muscle contraction regulates tendon development at the molecular, signaling, structural, and functional levels in *mdg* mutants. We find that without active muscle contraction, tendons are thinner due to incompetencies in both cell proliferation and matrix organization. Using transcriptomic analyses from single cell RNA-sequencing (scRNA-seq) and bulk RNA-sequencing (RNA-seq), we identify key matrix and cytoskeletal components that are downregulated in the absence of muscle loading. Furthermore, we demonstrate that modulation of TGFβ signaling is disrupted following muscle paralysis, and that the downregulation of TGFβ signaling is, in fact, necessary for normal tendon development. Finally, we show that the epitenon fails to form in muscle-paralyzed embryos. Collectively, these data establish a comprehensive analysis of the effects of embryonic muscle paralysis on tendon development and identify key signaling dynamics required for continued tendon growth.

## Results

### Muscle loading is required for the proliferation of embryonic tendon cells

Consistent with previous studies, the developing tendons in both the fore- and hindlimbs of *mdg* embryos were significantly thinner and shorter than their wild-type (WT) littermates at E18.5 (**Fig. 1A-H**)(*14*). *Mdg* Achilles tendons were consistently shorter than WT littermates, even when normalized to tibial length (**Fig. 1I**). This suggests that the size defects in *mdg* tendons are not secondary to a skeletal defect but rather reflect impaired tendon growth intrinsically. Since *mdg* tendons appeared grossly normal at E14.5, we evaluated tendon cell proliferation in Achilles tendons by EdU incorporation analyses from E14.5 to E16.5. As expected, proliferation was comparable in *mdg* and WT embryos at E14.5. However, by E15.5, *mdg* tendon cell proliferation declined from WT levels and remained low at E16.5 (**Fig. 1I-J**). The overall cellular density was comparable between WT and *mdg* tendons at all stages examined, suggesting that the lateral growth defects in the *mdg* tendons arise, in part, from the diminished proliferation of the resident tendon cells (**Fig 1K)**.

**Fig. 1.**
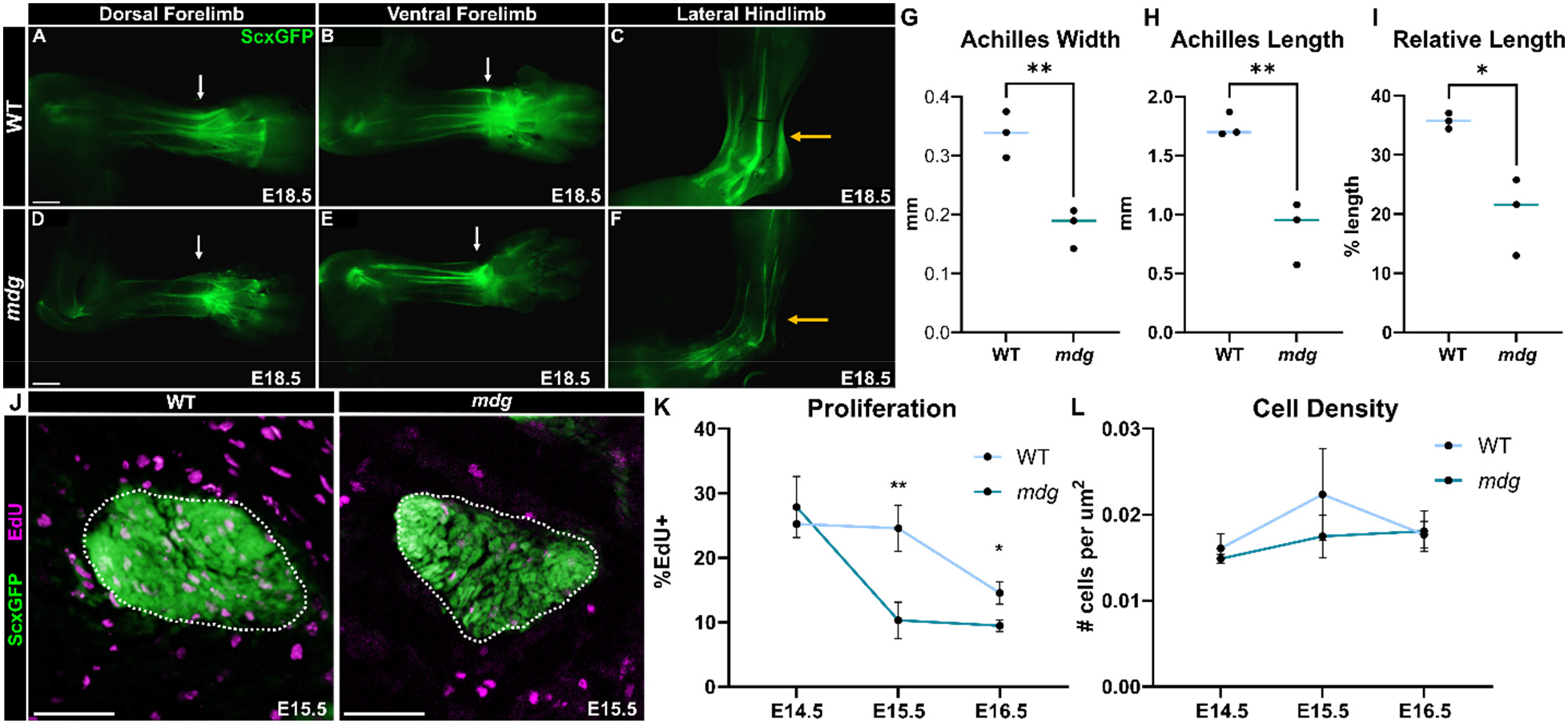
*Mdg* tendons are smaller and less proliferative. (**A-F**) Whole-mount images of *ScxGFP* in WT and *mdg* forelimbs (**A, B, D, E**) and hindlimbs (**C, F**) at E18.5. Arrows point to affected flexor and extensor tendons (white) and Achilles tendons (yellow). (**G, H**) Whole-mount quantification of Achilles tendon width and length from hindlimb images at E18.5 (n=3, two-tailed Student’s *t* test). (**I**) Achilles tendon length as a percent of limb length (calcaneus to tibial surface). (**J**) EdU detection in transverse midsubstance Achilles sections (ScxGFP: Cya, EdU: Magenta). White dashes denote Achilles tendon. (**K**) Quantification of EdU+ cells per nuclei E14.5-E16.5 (n=3, two-tailed Student’s *t* test, stars indicate significance between genotypes at each timepoint). (**L**) Quantification of cells per tendon area in transverse Achilles section E14.5-E16.5. Scale bar: 1mm (**A-F**), 50uM (**J**). For all quantifications, *p < 0.05,

### The developing tendon matrix is disrupted in the absence of muscle loading

To test the requirement for muscle loading in tendon matrix development, we examined the structural and functional properties of *mdg* mutants. Transmission electron microscopy (TEM) images of transverse sections through the Achilles tendon of E18.5 embryos revealed that *mdg* tendons have grossly larger collagen fibrils (WT: 17.03±2.32nm vs. *mdg:* 20.48±3.40nm, p<0.0001) (**Fig 2A-B**). Interestingly, although the *mdg* fibril sizes were larger, the fibrils were less densely packed within the tendon fascicle, suggesting impairment of normal matrix organization or alterations in the interfibrillar matrix (**Fig 2C**). This matrix disorganization was further reflected by second harmonic generation (SHG) imaging of longitudinal Achilles tendon sections (**Fig 2D**). There was significantly lower SHG signal throughout the *mdg* tendons. Since collagen fibrils are present in *mdg* tendons (**Fig. 2A**), the reduced collagen signal likely reflects disorganized fibril alignment or reduced packing density rather than a lack of collagen (**Fig. 2E**). Moreover, a reduction in phalloidin intensity within the *mdg* tendons suggests cytoskeletal disorganization (**Fig. 2D, E**). Functionally, *mdg* tendons showed an increased elastic modulus by AFM, likely attributable to larger collagen fibrils, collagen disorganization, or interfibrillar matrix alterations (**Fig. 1F**). There were no detectable differences in the viscoelastic properties, however, suggesting that energy dissipation in these tendons was not affected by the observed physical changes in collagen fibril or interfibrillar matrix (**Fig. 1G**). These results show that muscle loading is important for regulating the organization of the developing collagenous matrixand that matrix disorganization may contribute to functional deficits.

**Fig. 2.**
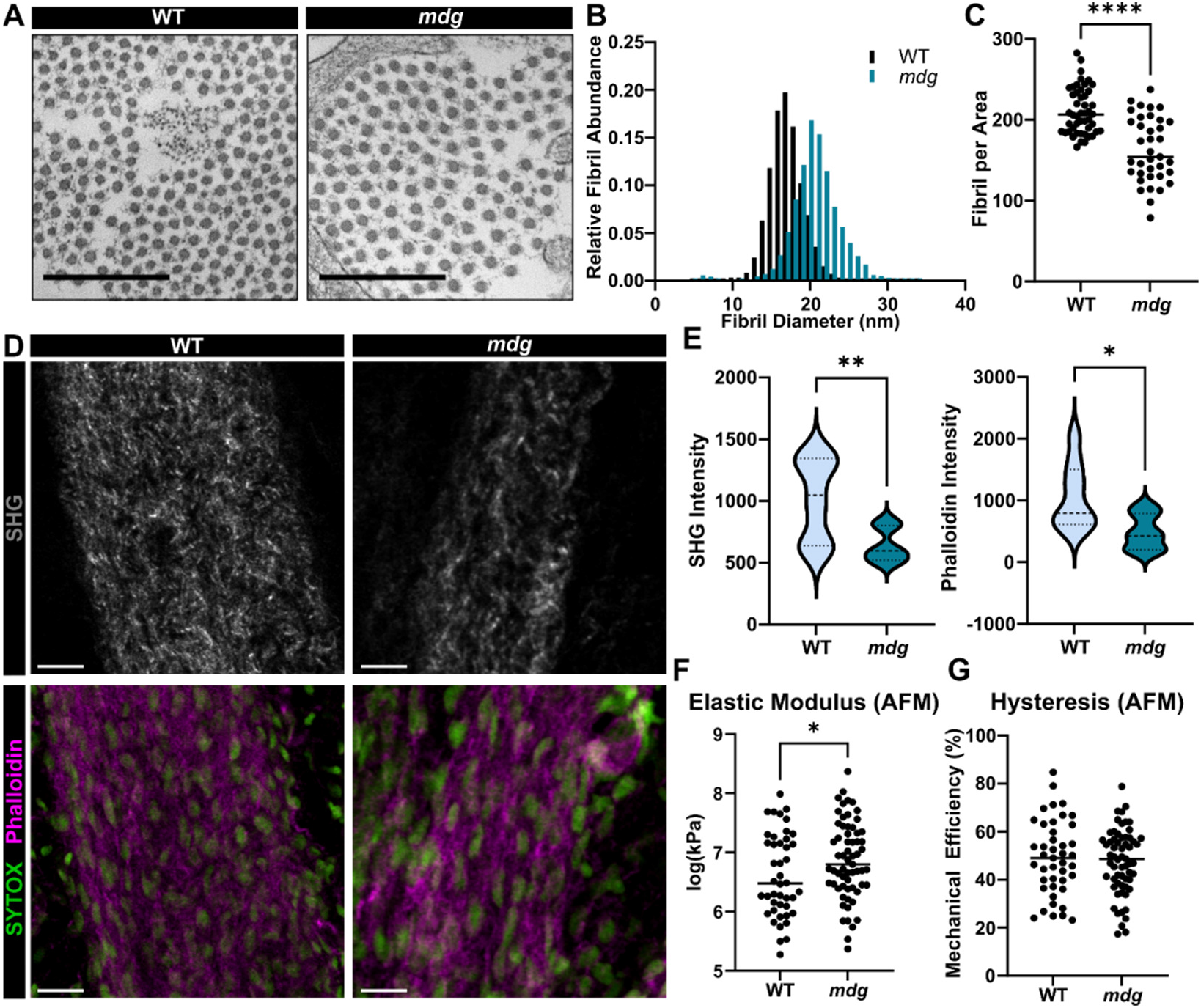
Matrix organization is inferior in *mdg* tendons. (**A**) Transmission electron microscopy images through E18.5 transverse Achilles sections. (**B**) Distribution of WT and *mdg* fibril size from TEM (**C**) Number of fibrils per measured area TEM (n=4 embryos, Kolmogorov-Smirnov test). (**D**) SHG and IHC of WT and *mdg* E18.5 longitudinal Achilles sections. (**E**) Quantification of SHG and Phalloidin intensity for representative regions of longitudinal sections (n-3, Kolmogorov-Smirnov test). (**F**). Elastic modulus and hysteresis from AFM on E18.5 longitudinal Achilles sections (n=6-8, two-tailed Student’s t test). For all quantifications, *p < 0.05, **p < 0.01, ****p < 0.0001. Scalebar: 500nm (**A**), 25um (**D**).

### Single cell RNA sequencing analyses reveal transcriptional dysregulation in mdg tendon cells

To identify molecular regulators of these functional deficits, we performed scRNA-seq on fore- and hindlimbs from E14.5 and E16.5 WT and *mdg* embryos (**Fig. 3A**). Initial coarse clustering using a low resolution (r=0.1) revealed 11 major cell clusters, based on their differentially expressed transcripts (**Fig. 3B,C**). Cluster 0 was identified as the major musculoskeletal cell population based on gene expression signatures for limb mesenchyme (*Prrx1*), cartilage (*Sox9*), and tendon (*Scx*) (**Fig. 3D**). E16.5 *mdg* cells contribute to ∼50% of the musculoskeletal cluster and were found to represent ∼25% to 50% of cells in all other clusters. Cells from all genotypes and timepoints are present in the identified clusters (**Fig. 3E,F**).

**Fig. 3.**
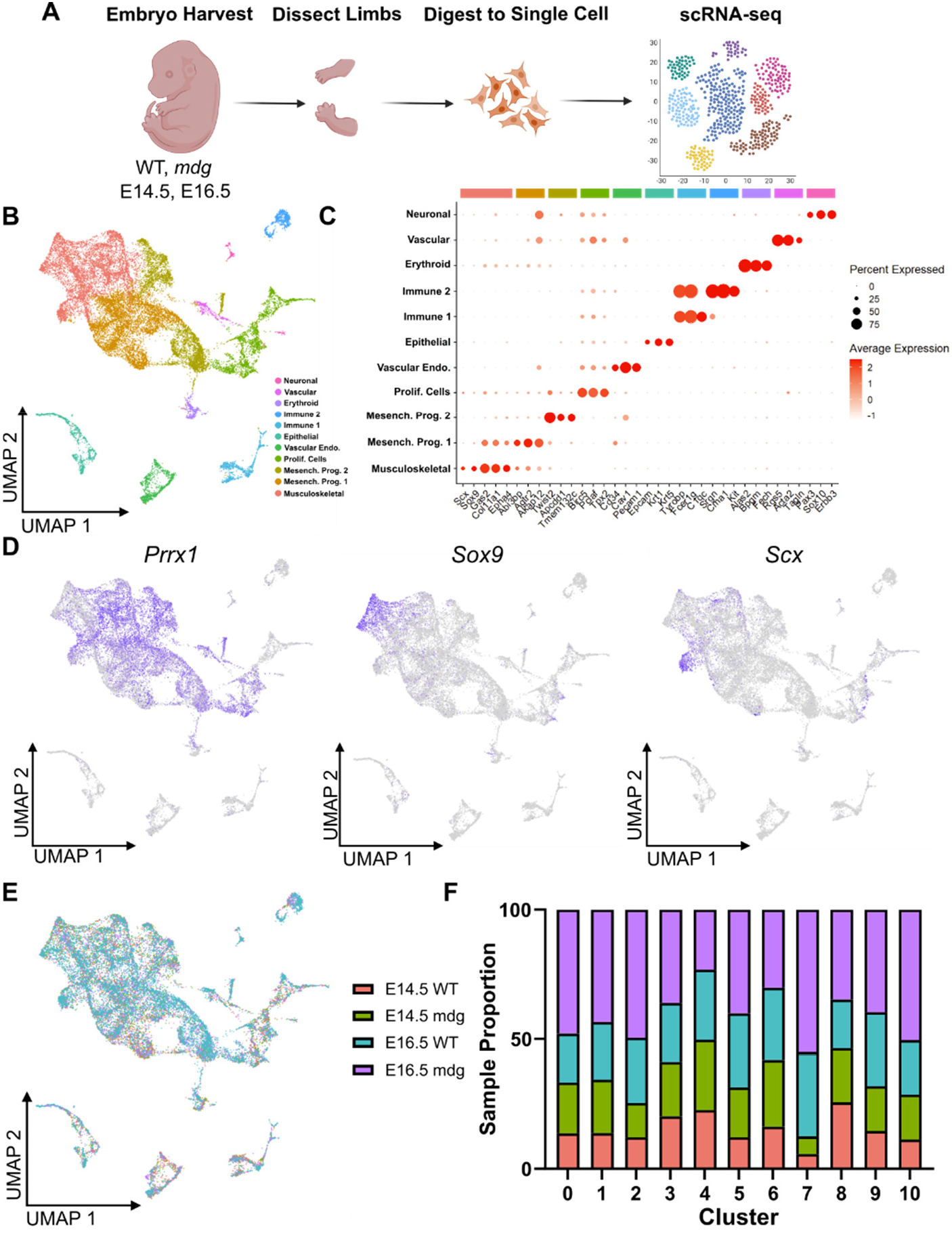
scRNA-seq reveals cell populations in developing limbs. (**A**) Schematic for scRNA-seq sample collection. (**B**) Unbiased clustering of limb cell scRNA-seq as UMAP. (**C**) Dot plot of top differential markers per cluster. (**D**) Expression plots in UMAP space of *Prrx1, Sox9*, and *Scx*. (**E, F**) Cell contributions to cluster by genotype and timepoint.

To understand how specific musculoskeletal populations are affected in the absence of muscle contraction, the musculoskeletal cluster was subset and re-embedded (**Fig. 4A**). The resulting 7 populations were identified based on their inter-cluster differentially expressed genes (DEGs) as: cartilage, skeletal progenitors, autopod and zeugopod progenitors, proximal and distal progenitors, connective tissue progenitors, tendon, and two dorsal and ventral progenitor clusters (**Fig 4B**). E16.5 *mdg* cells contribute the majority of cells to the tendon cluster, but cells from all timepoints and genotypes are represented in all clusters (**Fig 4C, D)**.

**Fig. 4.**
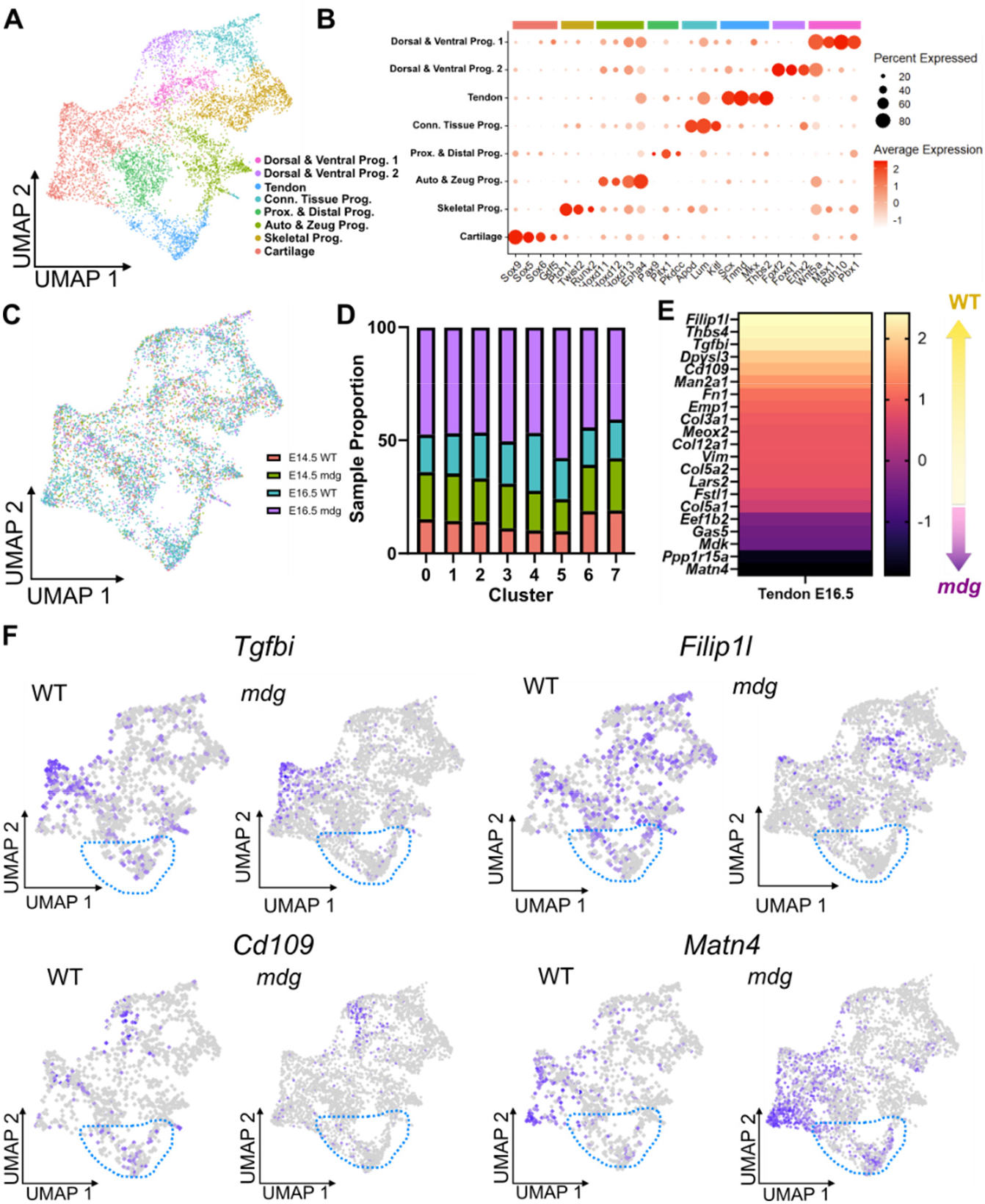
scRNA-seq reveals transcriptional differences between WT and *mdg* tendons at E16.5. (**A**) Unbiased clustering of re-embedded musculoskeletal cell cluster as UMAP projection. (**B**) Dot plot of cluster-defining differentially expressed genes. (**C, D**) Cell contributions to cluster split by genotype and timepoint. (**E**) Differentially expressed genes between WT and *mdg* tendon cluster for E16.5 cells (log2FC > 0.25, p.adj < 0.05). (**F**) Expression plots for *Tgfbi, Filip1l, Cd109*, and *Matn4* in UMAP space split by genotype. Blue lines denote tendon cluster.

DEG analyses using non-parametric Wilcoxon rank sum tests revealed no differentially expressed genes between WT and *mdg* tendon cells at E14.5. By E16.5, there were 21 DEGs between WT and *mdg* tendon cells (**Fig 4E, F**). Interestingly, expression of transcription factors associated with tendon cell fate (*Scx, Mkx*) were not different (**Fig 5A,E**). Many of the genes that were enriched in WT compared to *mdg* were related to tendon matrix components or matrix regulation (*Filip1l, Thbs4, Timp3, Col3a1, Col5a2, Col12a1, Vim*) and TGFβ signaling (*Tgfbi, Cd109*) (**Fig. 4E, F**). *In situ* hybridization (RNAScope) for selected targets – *Thbs4* (matrix organization), *Meox2* (mesenchymal transcription factor), and *Timp3* (matrix turnover) – confirmed significantly reduced expression in *mdg* tendons throughout the developing forelimb tendons (**Fig. 5B-E, G-K**). Collectively, these data suggest that once embryonic tendon cells adopt their tenogenic fate (prior to E14.5), they maintain their fate in the absence of muscle contraction. However, the development of the tendon matrix is disrupted, in part due to downregulated expression of matrix component and organization genes.

**Fig. 5.**
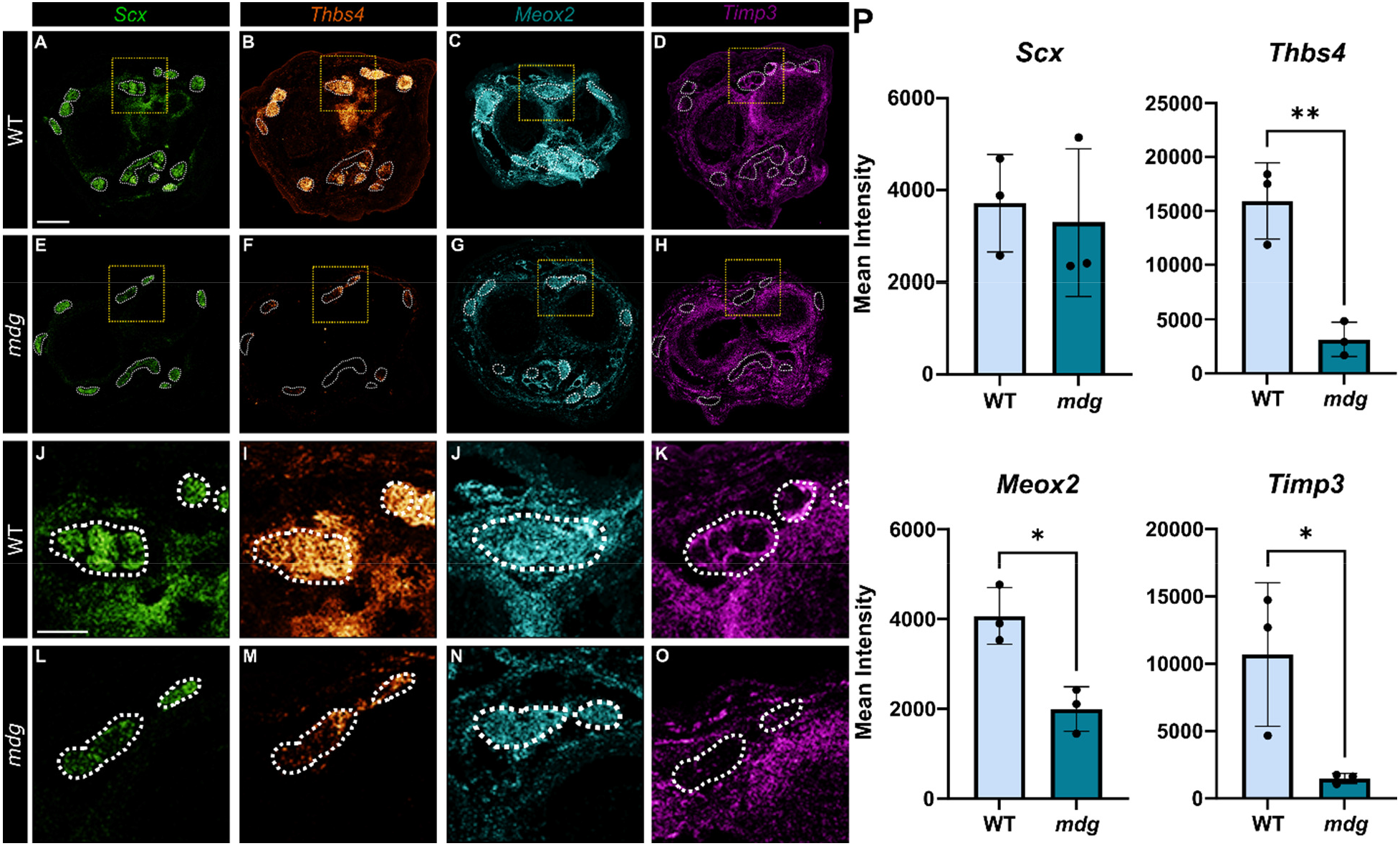
*In situ hybridization* confirms downregulation of DEGs. (**A-O**) E16.5 WT and *mdg* transverse sections through the wrist for: (**A, E, J, L**) *Scx;* (**B, F, I, M**) *Thbs4*; (**C, G, J, N**) *Meox2*; (**D, H, K, O**) *Timp3*. White dashes indicate tendons. (**P**) Quantification of ISH intensity in extensor tendons (n=3, two-tailed Student’s t test). For all quantifications, *p < 0.05, **p < 0.01. Scalebar: 200um (**A-H**), 50um (**J-O**).

### Tgfβ signaling is disrupted in the developing mdg tendons

Having established that matrix production and turnover is disrupted in the developing *mdg* tendons, we then investigated whether there is an intermediate signaling pathway altered with muscle paralysis that might be mediating these changes.

Pathway inferences analysis revealed that TGFβ signaling was specifically enriched in the tendon cell cluster, although this analysis did not discriminate between genotype and timepoint (**Fig. 6A**). Gene Set Enrichment Analysis (GSEA) between WT and *mdg* mutants primarily identified matrix and cytoskeletal pathways as differentially enriched between the two groups, but it likewise identified TGFβ related terms enriched in the data set, although in WT cells (**Fig 6B**). Moreover, at E16.5, two genes related to TGFβ signaling (*Tgfbi, Cd109*) were elevated in the wild-type tendon cell population compared to their *mdg* counterparts (**Fig. 4E**). *In situ hybridization* for *Tgfbi* confirmed reduced expression in the *mdg* limbs at E16.5 (**Fig. 6C, D**). At E14.5, both wild-type and *mdg* tendons had a high level of active TGFβ signaling, as ∼80% of tendon cells were positive for pSMAD2/3 at this stage. In wild-type embryos, this declined to only around ∼20% of tendon cells by E15.5, while *mdg* tendons retained a significantly higher percentage of positive cells (∼50%) (**Fig. 6E, F**). By E16.5, WT tendons have shut off TGFβ signaling, where only ∼5% of cells were still positive for pSMAD2/3. The *mdg* mutants, however, retained an elevated level of TGFβ signaling (∼50%), comparable to their E15.5 stage. This data suggests that the disruption of normal tendon growth due to muscle paralysis may be due to the failure of *mdg* tendon cells to turn off TGFβ signaling.

**Fig. 6.**
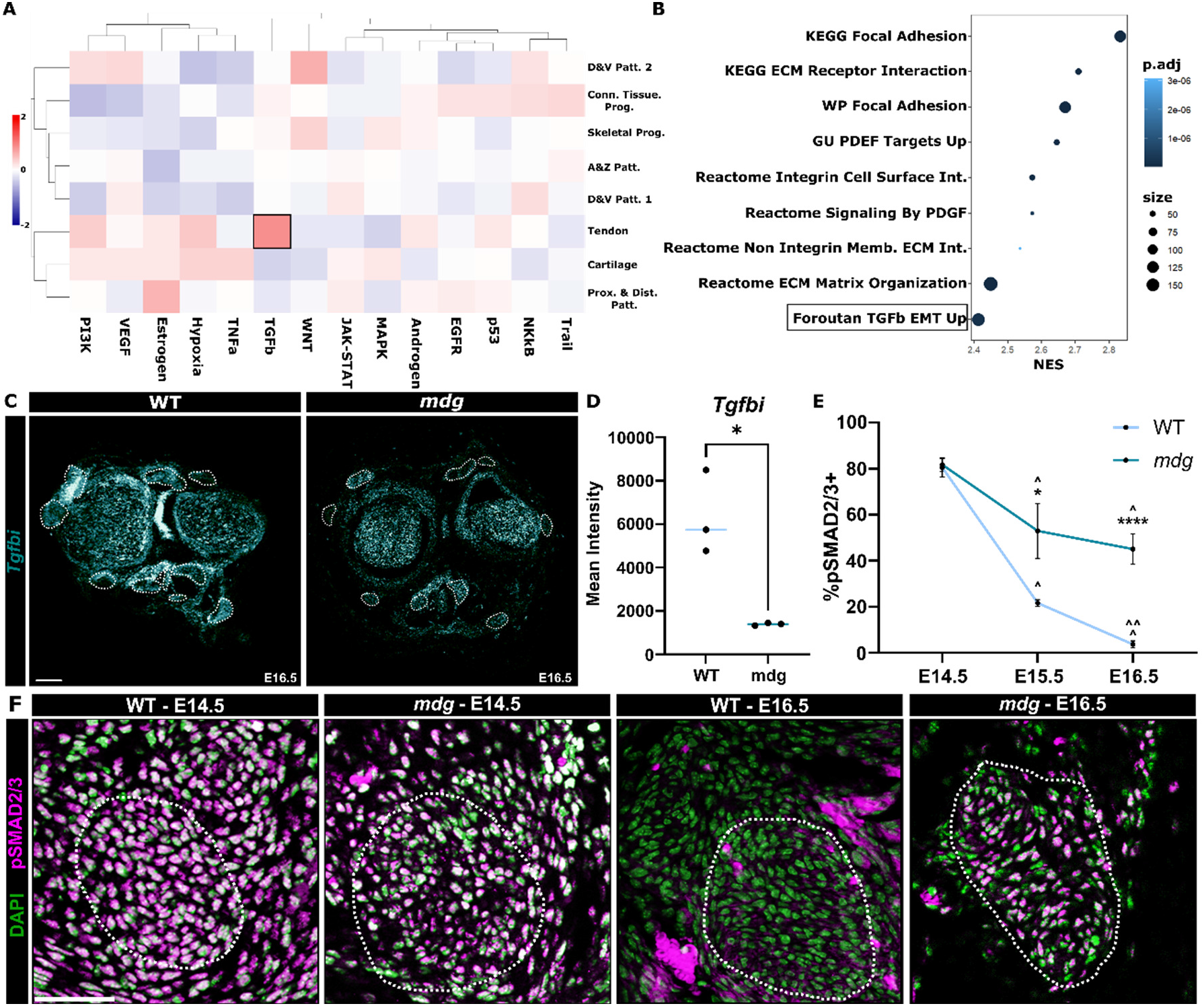
TGFβ signaling is dysregulated in *mdg* tendons. (**A**) Pathway inference analysis between clusters. Colors denote positive (red) or negative (blue) pathway enrichment score. Black box highlights TGFβ enrichment in tendon cell cluster. (**B**) GSEA enrichment plot of select enriched pathways in E16.5 tendon cluster on DEGs from WT and *mdg*. NES denotes Normalized Enrichment Score. (**C**) ISH for *Tgfbi* and *Cd109* on E16.5 WT and *mdg* transverse wrist sections. (**D**) Quantification of ISH intensity in *mdg* forelimb extensor tendons relative to WT (n=3, two-tailed Student’s t test). (**E**) Quantification of pSMAD2/3+ cells per nuclei (n=3, two-tailed Student’s t test, stars indicate significance between genotypes, carrots indicate significance by embryonic stages with a genotype vs. E14.5). (**F**) IHC for pSMAD2/3 in E14.5 and E16.5 WT and *mdg* tendons. Dotted white lines denote Achilles tendons as detected from serial sections by ScxGFP expression. For all quantifications, *p < 0.05, **p < 0.01, ****p < 0.0001. Scalebar: 100um (**A**), 50um (**F**).

### The developing epitenon does not form in the absence of muscle loading

While scRNA-seq permits cellular level transcriptomic analysis at these early stages, we hypothesized that some of the transcriptional differences may have been lost due to dropout events (*27*). To complement the scRNA-seq transcriptional analyses, we dissected fore- and hindlimb tendons from E18.5 WT and *mdg* embryos, isolated tendon RNA, and performed RNA-seq (**Fig. 7A)**. E18.5 WT and *mdg* tendons cluster closely in the principal component space, suggesting transcriptional similarity and supporting our E16.5 scRNA-seq findings that WT and *mdg* tendons retain many phenotypic features (**Fig. 7B**). Interestingly, we found 72 genes were upregulated and 21 genes were downregulated in the WT tendons compared to *mdg*, more than what we were able to detect in our single cell dataset (**Fig. 7C)**. Many of the genes enriched in the WT tendons were cytoskeletal or actomyosin components (*Pdlim1, Acta1, Myl2*), reflected by the enrichment of GSEA terms related to actomyosin and contractility (**Fig. 7D,E**).

**Fig. 7.**
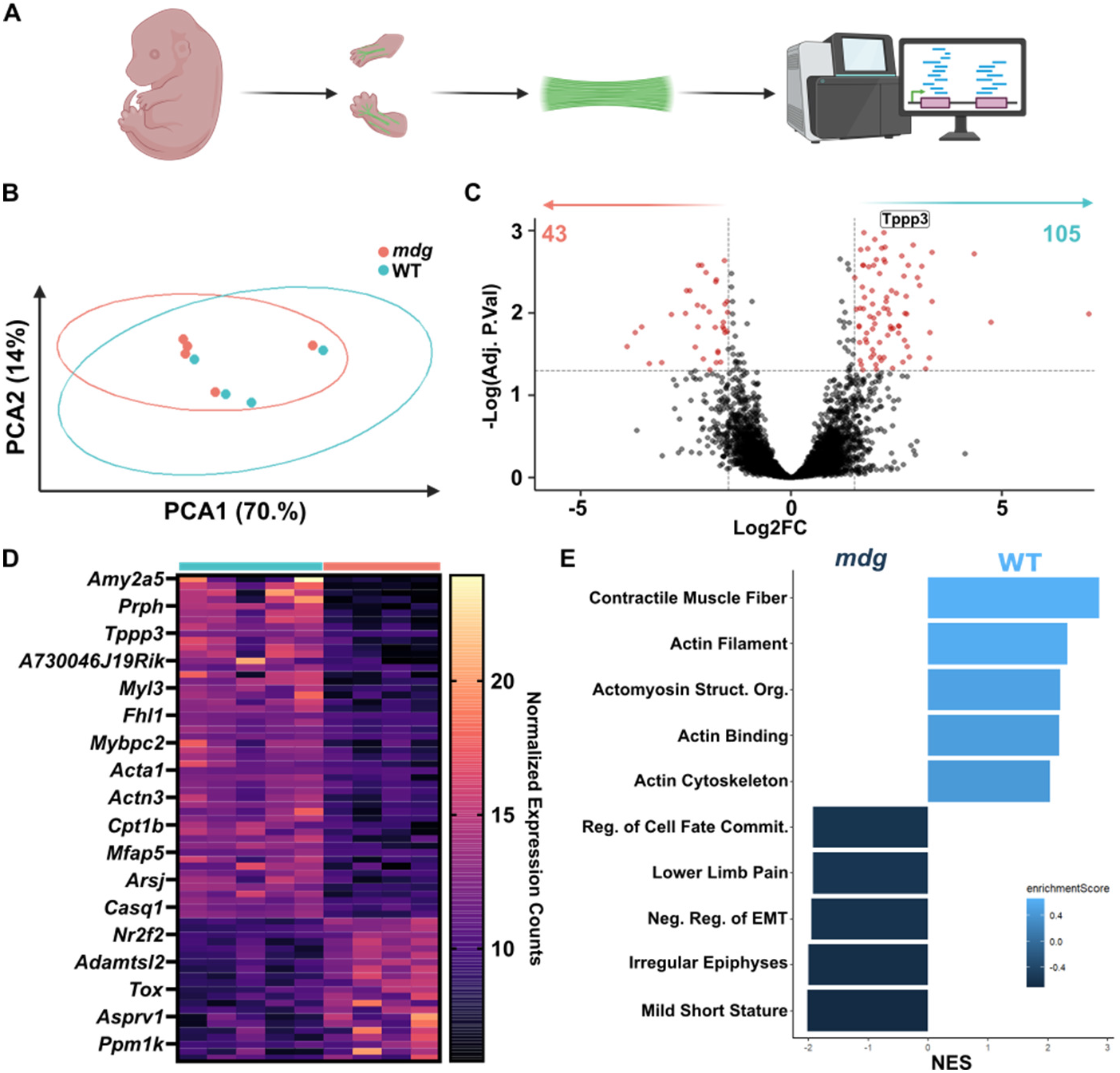
RNA-seq reveals actomyosin and cytoskeletal components are dysregulated in *mdg* tendons. (**A**) Schematic of sample collection for RNA-seq. (**B**) E18.5 WT and *mdg* transcriptomes in the PC space. (C-D) Volcano plot (**C**) and heatmap (**D**) of DEGs where positive Log2FC is enriched in WT and negative is enriched in *mdg* (DEGs: log2FC > 1.25, p.adj < 0.05). Heatmap value corresponds to normalized transcript expression. (**E**) Select enriched pathways in WT and *mdg* tendons from GSEA analysis, where NES is Normalized Enrichment Score (p.adj < 0.05).

Notably, *Tppp3*, a marker of the epitenon, is enriched in WT compared to *mdg* tendons (**Fig. 7C**). Taken together with the complete absence of *Timp3* epitenon expression in the *mdg* mutant at E16.5 (**Fig 5**), we tested the possibility that epitenon formation may be disrupted in *mdg* mutants. To determine whether there was an epitenon phenotype, we performed *in situs* for *Tppp3* throughout the forelimb of embryos from E13.5 to E16.5. Expression appeared grossly normal between E13.5 WT and *mdg* embryos with *Tppp3* expression primarily detected in the adjacent muscles (**Fig. 8B**). However, by E14.5, *Tppp3* expression around the tendons was completely lost in *mdg* mutants and remained absent at E16.5. Importantly, *mdg* mutants still showed clear expression of *Tppp3* in the developing muscles and around some joints (**Fig 8B**). This demonstrates that loss of *Tppp3* was a failure of epitenon formation, rather than an overall lack of *Tppp3* expression in *mdg* mutants. Collectively, this data suggests that signaling from muscle loading is required for the formation of the epitenon.

**Fig. 8.**
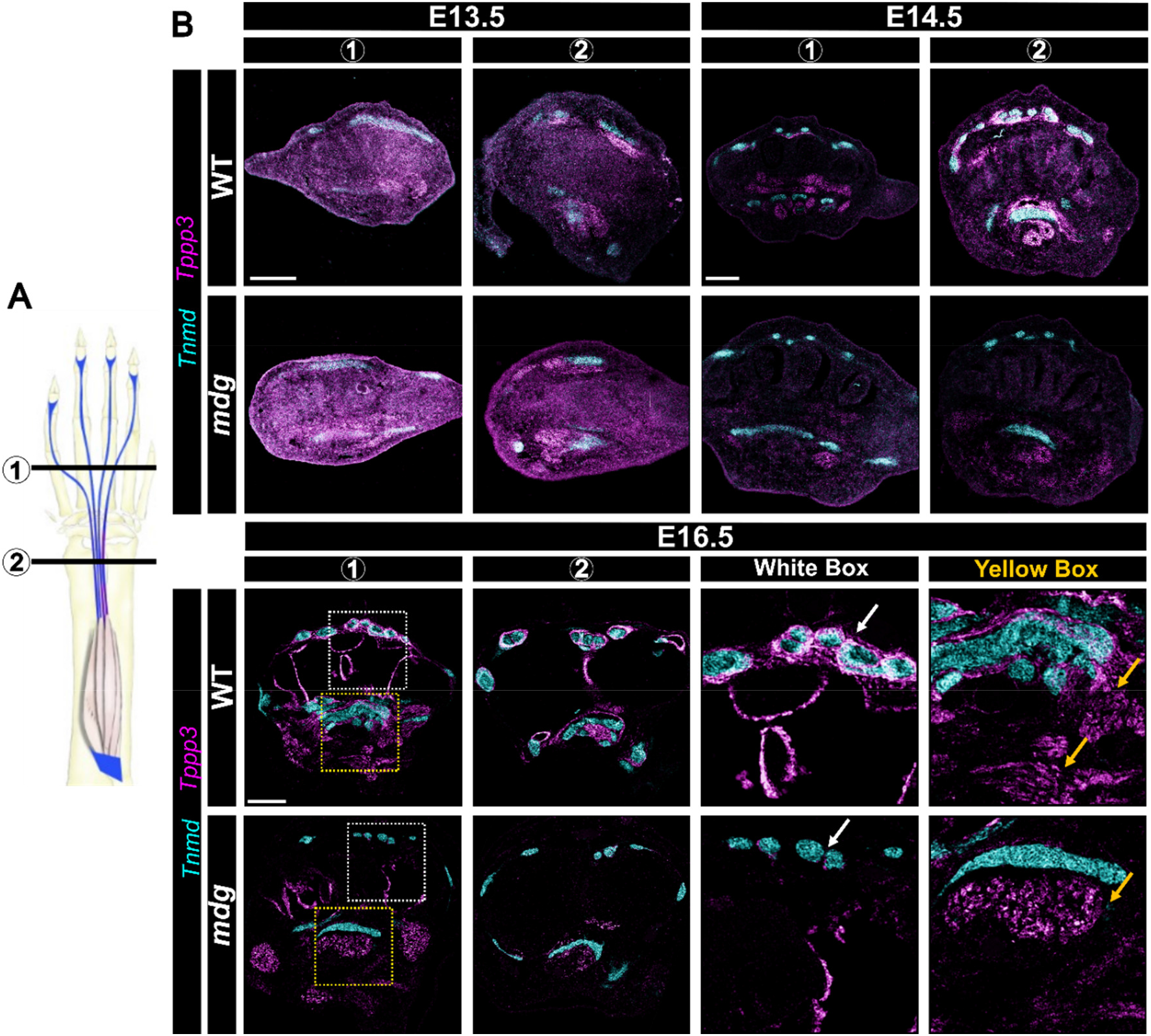
The epitenon does not form in *mdg* mutants. (**A**) Schematic of paw (1) and wrist (2) levels for ISH analysis. (**B**). ISH for *Tnmd* (cyan) and *Tppp3* (purple) at paw (1) and wrist (2) levels in E13.5, E14.5, and E16.5 in WT and *mdg* mutants. Arrows indicate tendons/epitenon (white) and muscle (yellow). Scale bar: 500uM (E13.5); 200uM (E14.5, E16.5).

## Discussion

In this study, we show that muscle contraction is a required cue for embryonic tendon development, regulating tendon cell proliferation, matrix deposition/organization, and functional properties (**Fig. 9**). Surprisingly, the maintenance of tendon cell identity (as defined by established transcription factors *Scx* and *Mkx*) was independent of muscle contraction. In addition, we found that attenuation of TGFβ signaling occurs during the tendon growth stage, and this was linked to muscle contraction as paralyzed *mdg* embryos maintained elevated of TGFβ signaling. Finally, we showed that mechanical signals are required for the formation of the epitenon, which is completely lost in *mdg* embryos.

**Fig. 9.**
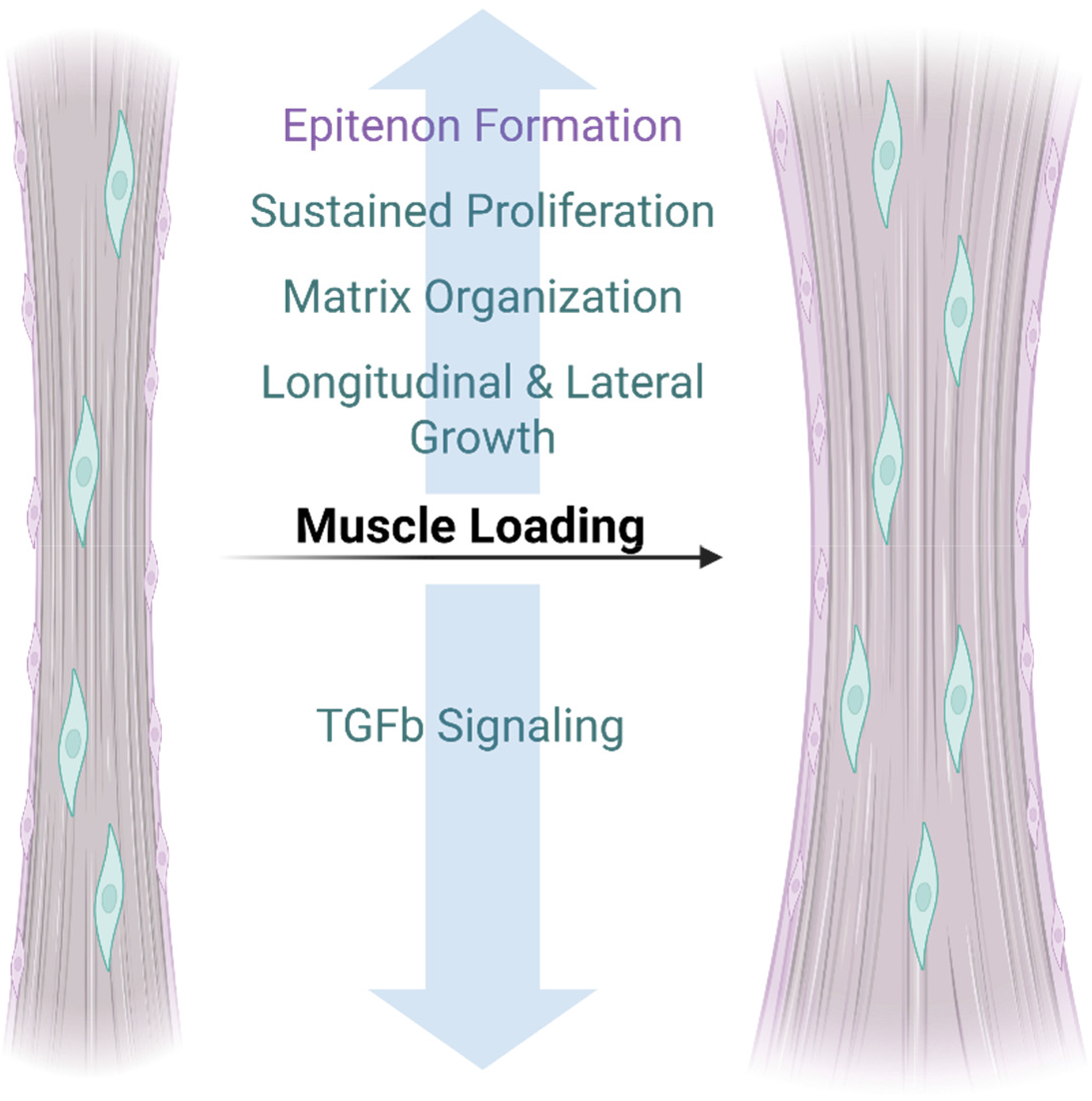
Muscle loading dependency in embryonic tendon development. Schematic of tendon embryonic events during the growth stage that require muscle loading.

### Mechanical forces from muscle loading are required for matrix production and organization in developing tendons

Our data support the general consensus that a loss of mechanical loading results in poor matrix organization in connective tissues (*26, 28*). Recent studies in chick have shown that elastic modulus is decreased with muscle paralysis with no changes in fibril packing (*18, 28*). Although we found that the indentation modulus in *mdg* tendons is higher relative to WT, this may be due to alterations in the matrix structure or organization. Another key difference is that the *mdg* mutation causes continuous muscle paralysis, whereas pharmacological paralysis may be targeting later stages of tendon maturation. Additionally, AFM conducted here consisted of compression in the plane normal to the longitudinal axis, not necessarily reflective of physiological tendon loading. We were limited to AFM due to the very small size of embryonic tendons, which precludes more traditional tensile mechanical testing. We interpret the higher modulus in *mdg* tendons to reflect the observed increase in fibril area, disorganization of collagen fibrils, or aberrant interfibrillar matrix or fiber cross-linking. In support of this hypothesis, we consistently see *Matrilin4 (Matn4)*, an ECM protein that can interact with collagens and promote the formation of filamentous structures (*29*), enriched in *mdg* tendons. *Matn4* enrichment in the *mdg* tendons may indicate increased networking between the collagen fibers, which is then reflected functionally in a higher compressive modulus.

From our transcriptional profiling of *mdg* tendons, we identified specific matrix components associated with differences in ECM composition and organization. *Thrombospondin4 (Thbs4)* encodes a matricellular protein that we and others identified as downregulated in tendon with muscle paralysis (Lipp et al., 2023a; Subramanian and Schilling, 2014; Subramanian et al., 2018). *Thbs4*^*-/-*^ mice have significantly larger collagen fibrils, similar to the phenotype we observe in *mdg* mutants (*33*). While *Thbs4* may be an important regulator of collagen fibril diameter, additional collagenous components (such as types III, V, and XII) in tendon are also important for regulating fibril size and organization, as well as cell-matrix organization (*34*–*36*). E16.5 *mdg* tendons showed reduced expression of *Col3a1, Col12a1, Col5a1*, and *Col5a2*. The increased fibril diameter and decreased fibril density observed in *mdg* tendons may therefore be attributed to the reduced expression of matrix components responsible for regulating collagen fibrillogenesis.

Transcriptional profiling at the E18.5 stage revealed several actomyosin and cytoskeletal components are dysregulated in *mdg* tendons. We identify more DEGs at this timepoint than from the single cell data. This may be attributable to the greater depth of sequencing with RNA-seq compared to scRNA-seq, or that the transcriptional differences are greater at later embryonic stages. The expression of actin and myosin components can be regulated downstream of mechanotransduction signaling pathways, such as MRTF/SRF and YAP/TAZ. One study has implicated dysregulated YAP signaling in chick tendons downstream of embryonic muscle paralysis, but this remains unexplored in embryonic mammalian models (*23*). In our RNA-seq data, none of the expected MRTF/SRF (*Acta2, Myl9, Vcl)* or YAP/TAZ *(Cyr61, Amotl1, Nuak2*) target genes were differentially expressed between WT and *mdg* tendons (*37*), suggesting that neither of these pathways are dysregulated with muscle paralysis. Future studies will incorporate epigenetic analyses to identify the upstream regulators of ECM and cytoskeletal genes in tendon development.

### Arrest of TGFβ signaling is regulated by mechanical signals during embryonic tendon growth

The elevated level of TGFβ signaling in E15.5 and E16.5 *mdg* tendons represents a clear link between mechanical loading from muscle and TGFβ signaling at this developmental stage. While TGFβ signaling is required for the induction of tendon progenitor cells at E12.5 (*38*), whether TGFβ is important for the subsequent differentiation and growth stages (E13.5 and E14.5+, respectively) previously remained unresolved. Targeted deletion of TGFβR2 with *ScxCre* did not show embryonic phenotypes, only early postnatal deterioration of tendons, suggesting TGFβ signaling is dispensable for tendons following progenitor induction until early postnatal events (*39*). Consistent with these findings, we find high levels of TGFβ signaling at E14.5 in WT embryos. Surprisingly, TGFβ signaling then declines dramatically during the growth stage (E15.5-16.5) in WT embryos but remains elevated in *mdg* tendons. These findings are intriguing, as they indicate that TGFβ is not merely dispensable at this stage, but that its attenuation is required for normal growth.

FGF and TGFβ signaling have previously been identified as two potential pathways regulating tendon development downstream of muscle loading in chick (Havis et al., 2016). One recent study showed that botulinum-toxin induced muscle paralysis of developing zebrafish reduced active TGFβ signaling in tendons (*32*). While we and others both link TGFβ downstream of muscle loading, we now find that muscle paralysis results in sustained, abnormal activation of TGFβ. While these findings seem contradictory, they likely point to the tightly coordinated timing of signaling events across distinct stages of tendon development (induction vs differentiation vs growth).

While we show that decreased activity of TGFβ signaling is essential for tendon growth, it remains unclear what other pathways may be responsible for positive regulation at this stage. One potential pathway is Wnt signaling as *Wnt2* is enriched in E18.5 WT tendons. In other tissue contexts, TGFβ and Wnt signaling pathways can share inhibitory or synergistic crosstalk (*40, 41*). Moreover, Wnt signaling can regulate the mechanoresponse of cells by modulating substrate rigidity, or alternatively be activated downstream of mechanical cues (*42, 43*). It is possible that muscle loading present during the tendon growth stage activates Wnt signaling which then directly or indirectly acts as an inhibitor of TGFβ. While Wnt signaling is important for early limb bud events and tendon-to-bone attachment, whether Wnt is important for embryonic tendon growth beyond this is unclear (*44–46*). Future studies are required to dissect the role, if any, of Wnt during this stage of tendon development.

### Formation of the epitenon requires mechanical loading from muscle

In recent years, numerous studies have identified the epitenon as a reservoir for postnatal stem cells that participate in tendon healing following injury (*47*–*49*). However, very little is known about the formation of the epitenon and what role it serves in tendon development. Interestingly, an early study of human embryonic development proposed that the differentiation of the tendon sheath tissue is a process that requires mechanical forces from adjacent tissue growth and muscle activity (*50, 51*). Our data now definitively show for the first time the necessity of muscle forces in this process.

While it is not clear whether the epitenon has any developmental role for tendon in the embryo, a few tendons that normally split at E15.5 remain fused in *mdg*. It is possible that *Timp3* secreted by epitenon is required for individuation of these tendons. Moreover, the more dispersed collagen fibrils in the *mdg* mutant may arise from a missing mechanical boundary around the tendon body in the absence of the epitenon. Previous studies have identified the formation of ectopic ligaments in the knee of *mdg* mutants. It is possible that the epitenon also serves as a barrier to control normal tendon and ligament patterning, and lack thereof may result in these observed ectopic tissues (*26*). An epitenon population was not captured in our scRNA-seq dataset for either timepoint or genotype. Additional study of the epitenon will be required to further understand why muscle paralysis prevents the formation of the epitenon, while having comparatively minimal effects on tendon cells.

Collectively, our results establish a profile of embryonic tendon cells in response to muscle paralysis. We reveal new roles for mechanical loading in epitenon formation and in the regulation of TGFβ signaling throughout tendon development. While our analyses focused on tendon cells to delineate their specific response to unloading, future studies on other affected cell types (such as the epitenon or connective tissue progenitors) will provide further insight into the intricately coordinated role of mechanical loading in musculoskeletal tissue development.

## Materials and Methods

### Mice

All animal work was approved by the Columbia University Institution for Animal Care and Use Committee. *Muscular dysgenesis (mdg*^*+/-*^) mice were crossed to *ScxGFP* mice to facilitate visualization of tendons(*52, 53*). For embryo collection, *mdg*^+/-^ males were time-mated with *mdg*^+/-^ females, with noon of the day a copulation plug was detected being defined as E0.5.

Heterozygous and null littermates served as controls for all assays and both are referred to in this manuscript as wild-type (WT).

### Immunohistochemistry and in situ hybridization

For *in situs* and immunofluorescence, collected embryos were fixed overnight in 4% paraformaldehyde (PFA) at 4°C. Following fixation, embryos were dehydrated serially in 5% and 30% sucrose and embedded in O.C.T. compound. 12μm cryosections of limbs were collected on slides for histological analyses. To quantify proliferation, pregnant dams were injected with 0.20mg EdU 2 hours prior to harvest. Proliferating cells were detected in transverse Achilles sections using ClickIT Plus EdU Imaging Kit (Invitrogen, C10634). For detection of RNA *in situ*, RNAScope Multiplex Fluorescent V2 (Advanced Cell Diagnostics) was carried out on transverse cryosections following the standard protocol. To detect Phospho-SMAD2/3, sections were incubated in high temperature sodium citrate for 10 minutes, followed by overnight incubation with Phospho-SMAD2/3 (Invitrogen, PA5-110155) and detected with AlexaFluor Plus 594 (Invitrogen, A-21207). IHC and ISH fluorescent images were acquired using Zeiss Apotome and processed in Zeiss Zen 3.8. Quantification was performed on transverse sections using ImageJ for a minimum of three embryos and two slices per embryo. Whole mount images were acquired using a Leica M165FC stereomicroscope with filters for fluorescence.

### RNA-sequencing

E18.5 WT and *mdg* embryos were dissected into ice-cold PBS. The fore- and hindlimbs were removed and skinned. Tendon dissection was guided by fluorescent visualization of *ScxGFP*. Dissected tendons were homogenized using TriZol, and RNA was isolated using the standard Trizol-Chloroform method (*54*). Isolated RNA was sequenced by Genewiz using their Ultra-Low Input pipeline. Raw reads were pseudoaligned to Mus musculus GRCm39 reference genome (Ensembl) for gene level counts. Differential gene expression analysis was performed using the standard pipeline for DESeq2 (*55*). Differentially expressed genes were defined as adjusted p.value < 0.01, and a log fold change of 1.5.

### Single Cell RNA-sequencing

E14.5 (n=5 WT, n=1 *mdg*) and E16.5 (n=5 WT, n=2 *mdg*) embryos were dissected into ice-cold PBS. Fore- and hindlimbs were dissected and digested to single cell at 37°C in 1.5mg/ml collagenase V (Worthington Biochemical, LS005280). Single-cell RNA-Seq was carried out by the Columbia Genome Center on an Illumina NovaSeq 6000 (10X Genomics, Single Cell 3’ v4).

Reads were aligned using cellranger (8.0.1) to the mouse genome (GRCm39-2024-A). Standard QC was performed to remove cells with abnormally high (>2500) and low (<500) unique reads. Analyses were performed using standard Seurat v5 integration analyses. Coarse clustering resolution (0.1) was used as a preliminary approach to identify major cell populations. The musculoskeletal cluster was identified as containing *Prrx1+, Scx+*, and *Sox9+* cells. This cluster was then subset and re-clustered (resolution 0.3) to refine the musculoskeletal populations. Clusters were identified through differential gene expression between clusters (p.adj<0.05). Differentially expressed genes between genotypes were identified by p.adj<0.05 and log2FC>0.25. Pathway inference was performed as described previously(*56*). Gene Set Enrichment Analysis (GSEA) was performed on differentially expressed genes using fgsea and MSigDB curated gene set collections(*57*).

### Transmission Electron Microscopy

Embryonic limbs were skinned and fixed in 1.5% paraformaldehyde/1.5% glutaraldehyde (Electron Microscopy Science) followed by post-fixation in 1% OsO4. Samples were then dehydrated in ethanol, embedded in Spurrs epoxy, and polymerized at 70°C over 18 hours. Four embryos were analyzed per genotype, with a minimum of 8 slices analyzed per embryo. Analysis was performed using a custom MATLAB scripts, described previously(*58*).

### Atomic Force Microscopy

Tendon indentation modulus was assessed using atomic force microscopy (Asylum MFP-3D, 6.1μm diameter polystyrene colloidal indenter, 0.02-0.77 N/m, 10nN indentation threshold, 1μm/sec indentation speed). Force-indentation curves were analyzed using built-in Hertz modeling within the Asylum software. Indentation modulus was natural log transformed as previously recommended to account for inherent spatial heterogeneity (*59*). Although the curve modeling methods commonly employed within AFM, including the Hertz model, assume an elastic half-space, inherent tissue viscoelasticity result in apparent curve nonlinearity and hysteresis. To capture potential differences in viscoelastic character, raw curves were also extracted and analyzed using a custom MATLAB code to assess mechanical efficiency, described as the percentage of conserved strain energy density between indentation and retraction. Statistical comparisons between genotype within tissue were conducted using a student’s t-test.

### Statistics

All quantitative data are presented as means ± SD. For IHC collected across multiple slices, technical replicates were averaged per biological replicate. For comparison of two groups, two-tailed unpaired Student’s t tests were performed. For comparison of distributions, Kolmogorov-Smirnov tests were performed. All statistical analysis was performed using GraphPad Prism v10.4.1. Sample sizes were determined based on our previous data, and no data were excluded.

## Acknowledgements

We thank Dr. Sarah Calve for providing the *mdg* mouse. We acknowledge Sara Tufa and Douglas Keene for TEM services and the Columbia Genome Center for their single cell sequencing services. We also thank Olga Yaringa for her contributions to this project. Graphical schematics were generated in BioRender.

## Author Contributions

Conceptualization: EK, AH

Data curation: EK, LC, AH

Formal analysis: EK, LC, AH

Funding acquisition: EK, AH, NC

Investigation: EK, LC

Methodology: EK, LC, NC, JS, AH

Project administration: EK, AH

Resources: LC, NC, AH

Software: EK, LC

Supervision: NC, JS, AH

Validation: EK, LC, AH

Visualization: EK

Writing – original draft: EK

Writing – review and editing: EK, LC, NC, JS, AH

## Data Availability

All data needed to evaluate the conclusions in the paper are present in the paper and/or the Supplementary Materials. All sequencing datasets will be uploaded to the Gene Expression Omnibus repository.

